# Sex-biased expression of enteroendocrine cell-derived hormones contributes to higher fat storage in *Drosophila* females

**DOI:** 10.1101/2025.09.09.675263

**Authors:** Puja Biswas, Elizabeth J. Rideout

**Affiliations:** Department of Cellular and Physiological Sciences, Life Sciences Institute, The University of British Columbia, Vancouver, Canada; Department of Pediatrics, BC Children’s Hospital Research Institute, The University of British Columbia, Vancouver, Canada

**Keywords:** Sex difference, gut, enteroendocrine cell, physiology, triglyceride, hormones, fat storage

## Abstract

Enteroendocrine (EE) cells in the *Drosophila* gut produce and release multiple factors, including Allatostatin A (AstA), Allatostatin C (AstC), neuropeptide F (NPF), tachykinin (Tk), Diuretic hormone 31 (Dh31), Bursicon, CCHamide 1, CCHamide 2, and short neuropeptide-F. Collectively, these peptides ensure that physiology (e.g., fat storage, fluid balance) and behavior (e.g., feeding, sleep) are coordinated with environmental factors such as nutrient quantity and quality. Despite notable sex differences in physiology and behavior, it remains unclear whether the regulation and function of these EE cell-derived factors are shared between males and females. Given that recent data identified sex-biased physiological effects of two EE cell-derived hormones on *Drosophila* food intake and energy mobilization, we performed a detailed characterization of these hormones in male and female flies. Despite an overall male bias in mRNA levels of *AstA*, *AstC*, *Tk*, *NPF*, *Dh31* in whole-body and head samples, we observed a strong female bias in mRNA levels of *AstC*, *Tk*, and *NPF* in the gut. To determine whether this sex-biased regulation was physiologically significant, we monitored triglyceride levels in flies with gut-specific knock-down of EE cell-derived hormones. In 5-day-old flies, knock-down of EE cell-derived AstC significantly reduced fat storage in females with no effect in males, whereas knock-down of EE cell-derived Tk produced a non-significant trend toward reduced fat storage in females. These female-specific effects on fat storage were reproduced in flies with neuron-specific knock-down of the AstC (AstC-R2) and Tk receptors (TkR99D). Together, these data uncover strongly sex-biased regulation of EE cell-derived hormones, and show that gut-specific knock-down of at least one of these hormones had a female-specific effect on body fat.

**Highlights:**

- Enteroendocrine cell-expressed hormones show strongly sex-biased expression
- Knock-down of enteroendocrine cell-derived AstC reduced body fat only in females
- Neuronal knock-down of AstC or Tk receptors reduced stored fat only in females

## 1. Introduction

In *Drosophila*, females store more fat than males [1–7]. Greater female fat storage has been observed in both mated and unmated females compared with age-matched males [1,3–7]. In flies, as in other animals, triglyceride is the main form of stored fat. Although triglyceride is present in many cell types and organs (*e.g.*, oenocytes, glia, neurons, gut) [5,6,8–14], the majority of triglyceride is stored in an organ called the fat body [3,12,15].

In females, high levels of triglyceride in the fat body play a key role in supporting reproduction and physiology [5,16]. Indeed, triglyceride from the fat body is a key energy source for the developing embryo [17]. Increased fat storage in adult females also supports prolonged survival during nutrient deprivation compared with males [4,6]. Despite these clear benefits of greater fat storage for female fertility, excess fat accumulation in males adversely affects their reproductive output. Specifically, males carrying a mutation that promotes excess triglyceride accumulation show reduced testis size, defects consistent with delays in spermatogenesis, and ultimately a reduction in sperm number [8]. The sex difference in fat storage therefore likely reflects the fact that males and females differ in how much whole-body triglyceride storage supports optimal fertility.

Recent studies have identified genes and pathways that contribute to sex differences in fat storage [5–7,18]. In mated females, the steroid hormone ecdysone acts on neurons to promote food intake, which is associated with increased body fat [5]. In unmated adult females, the insulin/insulin-like growth factor signaling pathway (IIS) plays a key role in maintaining an elevated level of triglyceride storage compared with adult males [18]. Specifically, adult females have higher production of *Drosophila* insulin-like peptide 3 (Dilp3) and greater insulin sensitivity, leading to higher IIS activity. This elevated IIS activity is important for females to store more triglyceride than males, as adult-specific ablation of the insulin-producing cells reduces body fat in females but not males [18].

In males, body fat levels are maintained by higher expression and activity of two catabolic pathways that promote fat breakdown. One pathway is regulated by triglyceride lipase *brummer* (*bmm*), where males show higher *bmm* mRNA levels compared with females [6]. This elevated *bmm* expression contributes to the sex difference in fat storage by restricting triglyceride accumulation in males. Similarly, males have higher production and secretion of Adipokinetic hormone (Akh), a key lipolytic hormone in insects [7]. As with *bmm*, high levels of Akh in males contribute to the sex difference in fat storage by limiting fat accumulation in males [7]. Together, these studies have defined a model of the sex difference in fat storage in which females maintain higher levels of fat storage in part due to a higher relative activity level for anabolic pathway IIS, whereas males have lower fat storage due to higher relative activity of catabolic effectors such as *bmm* and Akh. While some progress has been made in revealing the mechanisms underlying the sex-specific regulation of Akh and IIS [7,18], we do not have a complete understanding of the factors that determine the sex-biased regulation and function of these key metabolic factors.

Recent clues into the regulation of Akh, *bmm*, and IIS have emerged from studies on peptide hormone function [19–30], as several hormones influence physiology via effects on Akh- and Dilp-producing cells [19–22,24–33]. In particular, recent studies have illuminated an important role for hormones produced by the enteroendocrine (EE) cells of the gut [32,34–36]. Adult *Drosophila* EE cells produce and release hormones such as Allatostatin A (AstA), Allatostatin C (AstC), neuropeptide F (NPF), tachykinin (Tk), Diuretic hormone 31 (Dh31), Bursicon, CCHamide 1 (CCHa1) and CCHamide 2 (CCHa2), and short neuropeptide F (sNPF) [37–41].

EE cells are identified by expression of the homeodomain protein Prospero in adults [42,43], where EE cells that produce distinct hormones are present in anatomically defined regions of the adult gut [43]. For example, AstA- and Dh31-producing EE cells are located in the posterior midgut, whereas EE cells that produce AstC and Tk are found along the entire length of the midgut [37]. NPF-producing cells are found in the anterior and middle midgut [41]. Supporting a role for EE cell-derived hormones in regulating Akh/IIS, in fed conditions studies show EE cell-derived hormones such as Bursicon inhibit Akh secretion [28], whereas NPF enhances Dilp secretion from the insulin-producing cells [29]. During starvation, AstC promotes Akh release from Akh-producing cells to enable lipid mobilization [22]. While some EE cell-derived hormones have been shown to have sex-biased effects on food intake and energy mobilization [22,44,45], sex differences in the regulation and function of most of these hormones remain unclear. Defining potential differences in EE cells is an important task, as prior studies have revealed profound differences in gut biology between males and females.

For example, males and females differ in overall gut size and shape [46,47] and in the number of intestinal stem cell divisions [46,48–50]. The absorptive lining of the gut also shows sex differences during aging [49,51]. After mating, females show pronounced changes to gut size, function, and gene expression [46,52–55]. While sex differences in intestinal stem cells and enterocytes play a key role in mediating these differences in gut biology, we know less about male-female differences in EE cells. We therefore aimed to perform a detailed characterization of EE cell-derived hormones in males and females, and to determine the contribution of these hormones to physiology in each sex. We reveal profound sex-biased regulation of EE cell-expressed hormones AstC, Tk, and NPF: females show higher mRNA levels of these hormones in the gut than males. For at least one EE cell-derived hormone, this sex-biased regulation was physiologically significant, as knock-down of AstC in the gut reduced fat storage in females with no effect in males. Knock-down of Tk in the gut produced a trend toward reduced female fat storage. Female-specific fat storage defects were also observed following knock-down of AstC and Tk receptors in neurons and/or neuropeptide-producing cells. While the specific cell type targeted by these EE cell-derived hormones to influence fat storage remains unclear, this reveals a female-specific contribution of one EE cell-derived hormone in regulating body fat.

## 2. RESULTS

### 2.1. Sex differences in expression of gut-derived peptide hormones

Given that several EE cell-derived hormones such as AstA (FBgn0015591), AstC (FBgn0032336), Tk (FBgn0037976), NPF (FBgn0027109), Dh31 (FBgn0032048) are expressed in cells outside the gut [20,56–70], we used quantitative real-time PCR (qPCR) to assess whole-body mRNA levels of genes encoding EE cell-expressed hormones. In particular, we focused on hormones known to influence whole-body fat metabolism [20,22,29,44,71,72]. In 5-day-old *w^1118^* unmated adult males and females, we found that whole-body mRNA levels of *AstA*, *AstC*, *Tk*, *NPF*, and *Dh31* were significantly higher in males than in females (Figure 1A-E). To gain further insight into this sex-biased expression, we analyzed mRNA levels of these factors from isolated heads and intestines, as these are the main sites of AstA, AstC, Tk, NPF, and Dh31 production [35,37]. A significant male bias in mRNA levels was found in the head for *AstA*, *AstC*, *Tk*, *NPF,* and *Dh31* (Figure 1F-1J). In contrast, mRNA levels of *AstC*, *Tk*, and *NPF* in isolated intestines showed a strong female bias (Figure 1L-1N). No sex bias in the expression of *AstA* or *Dh31* was observed in the gut (Figure 1K, 1O).

**Figure 1.**
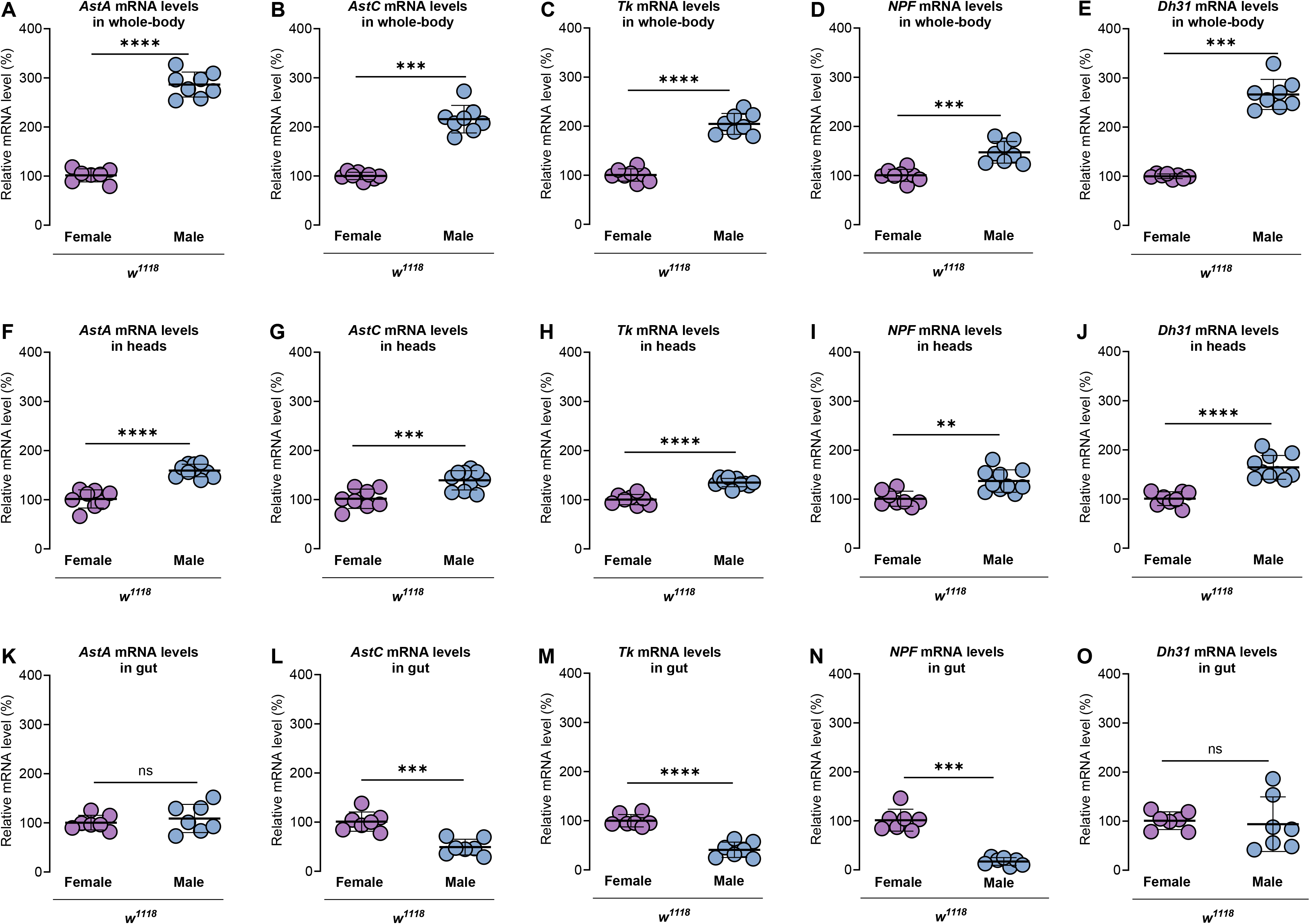
Sex differences in expression of gut-derived peptide hormones. (A-E) mRNA levels of *AstA* (*p*<0.0001; Student’s *t*-test) (A), *AstC* (*p*=0.0002; Mann-Whitney test) (B), *Tk* (*p*<0.0001; Student’s *t*-test) (C), *NPF* (*p*=0.0001; Student’s *t*-test) (D), *Dh31* (*p*=0.0002; Mann-Whitney test) (E) in whole-body were significantly higher in 5-day-old *w^1118^* males compared to females. *n*=7-8 biological replicates. (F-J) mRNA levels of *AstA* (*p*<0.0001; Student’s *t*-test) (F), *AstC* (*p*=0.001; Student’s *t*-test) (G), *Tk* (*p*<0.0001; Student’s *t*-test) (H), *NPF* (*p*=0.0015; Student’s *t*-test) (I), *Dh31* (*p*<0.0001; Student’s *t*-test) (J) in heads were significantly higher in 5-day-old *w^1118^* males compared to females. *n*=8-10 biological replicates. (K) mRNA levels of *AstA* (*p*=0.5039; Student’s *t*-test) in guts were not significantly different between 5-day-old *w^1118^*females and males. *n*=7 biological replicates. (L-N) mRNA levels of *AstC* (*p*=0.0002; Student’s *t*-test) (L), *Tk* (*p*<0.0001; Student’s *t*-test) (M), *NPF* (*p*=0.0006; Mann-Whitney test) (N) in guts were significantly higher in 5-day-old *w^1118^* females compared to males. *n*=7 biological replicates. (O) mRNA levels of *Dh31* (*p*=0.7517; Student’s *t*-test) in guts were not significantly different between 5-day-old *w^1118^*females and males. *n*=7 biological replicates. All data plotted as mean ± SEM. ns indicates not significant with *p*>0.05; ** *p*<0.01, *** *p*<0.001, **** *p*<0.0001. See also Figure S1.

Building on the sex bias in mRNA levels, we next examined mRNA levels of receptors that correspond to EE cell-expressed hormones with sex-biased expression in whole-body, fat body, and head samples. Whole-body mRNA levels of the receptors for AstA (*AstA-R2*), AstC (*AstC-R2*), Tk (*TkR99D*), NPF (*NPFR*), and Dh31 (*Dh31-R*) were significantly higher in 5-day-old *w^1118^*males compared with age-matched females (Figure 1-figure supplement 1A-E). For most peptides, the male bias was due to a higher mRNA level in the head and not the fat body (Figure 1-figure supplement 1A-E); however, *TkR99D* mRNA levels were higher in male fat bodies with no difference in head mRNA levels (Figure 1-figure supplement 1C). We therefore cannot rule out a contribution of additional anatomical sites to the male bias in expression of EE cell-expressed hormones, which is an interesting area for future investigation. Taken together with our data on peptide mRNA levels, our data suggest sex differences exist in both the expression of EE cell-derived hormones and in the ability of tissues to respond to available peptide.

### 2.2. Sex determination gene *transformer* does not regulate sex differences in EE cell-derived peptide mRNA levels

To determine the mechanism by which these differences in mRNA levels are established, we tested a role for sex determination gene *transformer* (*tra*). Normally, a functional Tra protein is only expressed in females, where Tra specifies most aspects of female sexual development and behavior [73–76]. Indeed, ectopic Tra expression in males is sufficient to specify many aspects of female sexual development and physiology [7,48,49,74,77,78]. Because *tra* mRNA is detected in the gut, specifically in ISC and EE cells [48,79], we asked whether broad overexpression of Tra in neurons and/or EE cells contributes to the sex difference in mRNA levels of EE cell-expressed hormones. For these data, cell type-specific Tra overexpression was considered to have a significant effect on EE cell-expressed hormones only if the experimental genotype (e.g., *tissue-GAL4>UAS-tra^F^*) significantly differed from both parental strains (e.g., *tissue-GAL4>+* and *+>UAS-tra^F^*) with the same direction of effect. We found that sex differences in mRNA levels of *AstA*, *AstC*, *Tk*, *NPF,* and *Dh31* were unaffected when we used either *voila-GAL4* (Figure 2A-2J) which expresses in EE and sensory cells, or *elav-GAL4* (Figure 2K-2T) which expresses in neurons and neuropeptide-producing cells, to drive Tra expression in these cells. However, we note that Tra expression in EE cells further augments the male bias in head *Tk* mRNA levels (Figure 2H), whereas Tra expression in female neurons paradoxically decreases *NPF* mRNA levels in the head (Figure 2S). Tra expression in neurons similarly had no effect on mRNA levels of *AstC-R2*, *TkR99D*, or *Dh31-R* in the head (Figure 2-figure supplement 2A-C). Thus, sex determination *tra* does not establish sex differences in levels of the mRNAs that encode either EE cell-derived hormones or their receptors.

**Figure 2.**
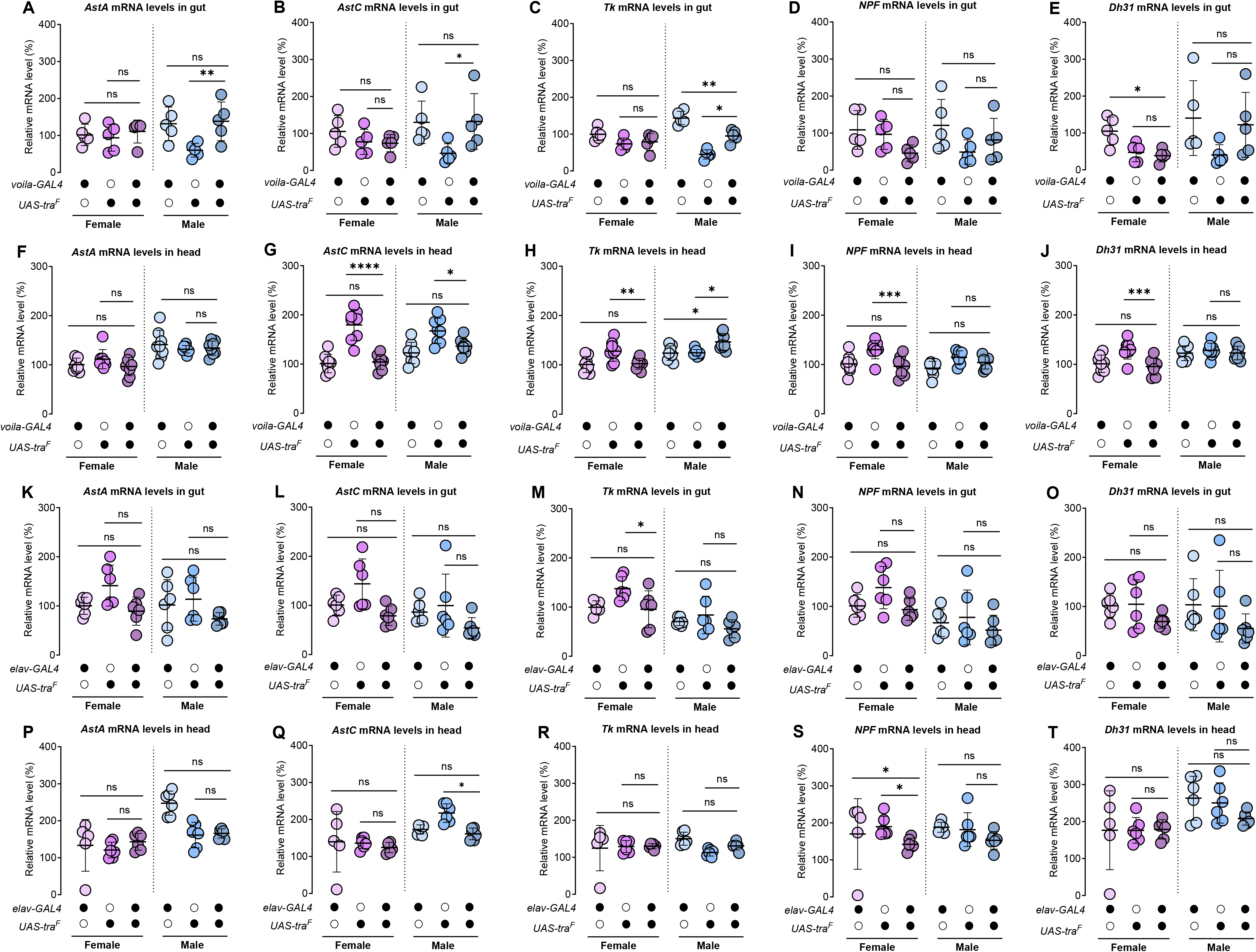
Sex determination gene *transformer* does not regulate sex differences in EE cell-derived peptide mRNA levels. For all data, cell type-specific RNAi was considered to have a significant effect only if the experimental genotype (e.g., *tissue-GAL4>UAS-RNAi*) significantly differed from both parental strains (e.g., *tissue-GAL4>+* and *+>UAS-RNAi*) with the same direction of effect. (A-E) mRNA levels of *AstA* (A), *AstC* (B), *Tk* (C), *NPF* (D), and *Dh31* (E) in the gut were measured in *voila-GAL4>UAS-tra^F^* flies and respective genetic controls (*voila-GAL4>+* and *+>UAS-tra^F^*) in females and males. Tra expression did not alter *AstA* (female: *p*^GAL4^>0.9999 and *p*^UAS^>0.9999; male: *p*^GAL4^>0.9999 and *p*^UAS^=0.0084), *AstC* (female: *p*^GAL4^=0.6814 and *p*^UAS^=1.0; male: *p*^GAL4^=0.9965 and *p*^UAS^=0.0463), *Tk* (female: *p*^GAL4^=0.2258 and *p*^UAS^>0.9999; male: *p*^GAL4^=0.0006 and *p*^UAS^=0.0004), *NPF* (female: *p*^GAL4^=0.1579 and *p*^UAS^=0.3389; male: *p*^GAL4^=0.6639 and *p*^UAS^=0.9043) or *Dh31* (female: *p*^GAL4^=0.0439 and *p*^UAS^=0.9745; male: *p*^GAL4^=0.9953 and *p*^UAS^=0.1370) levels in either sex. Sex:genotype interaction: *AstA* (*p*<0.0001), *AstC* (*p*=0.1078), *Tk* (*p*=0.0004), *NPF* (*p*=0.1655), *Dh31* (*p*=0.1945). Data were analyzed by two-way ANOVA with Bonferroni or Tukey’s HSD post-hoc tests as appropriate (aligned rank transform applied for non-parametric data in B and E); *n*=5 biological replicates. (F–J) mRNA levels of *AstA* (F), *AstC* (G), *Tk* (H), *NPF* (I), and *Dh31* (J) in the head were measured in *voila-GAL4>UAS-tra^F^* flies and respective genetic controls (*voila-GAL4>+* and *+>UAS-tra^F^*) in females and males. Tra expression did not alter *AstA* (female: *p*^GAL4^>0.9999 and *p*^UAS^=0.3344; male: *p*^GAL4^>0.9999 and *p*^UAS^>0.9999), *AstC* (female: *p*^GAL4^>0.9999 and *p*^UAS^<0.0001; male: *p*^GAL4^=0.6687 and *p*^UAS^=0.0236), *NPF* (female: *p*^GAL4^>0.9999 and *p*^UAS^=0.0006; male: *p*^GAL4^=0.5030 and *p*^UAS^=0.6158) or *Dh31* (female: *p*^GAL4^>0.9999 and *p*^UAS^=0.0003; male: *p*^GAL4^>0.9999 and *p*^UAS^>0.9999) levels in either sex. In females, *Tk* (*p*^GAL4^>0.9999 and *p*^UAS^=0.0044) levels did not alter, but in males, *Tk* was increased compared with both controls (*p*^GAL4^=0.0120 and *p*^UAS^=0.0156). Sex:genotype interaction: *AstA* (*p*=0.2471), *AstC* (*p*=0.0189), *Tk* (*p*=0.0003), *NPF* (*p*=0.1408). *Dh31* (*p*=0.0403). Data were analyzed by two-way ANOVA with Bonferroni post-hoc tests; *n*=8 biological replicates. (K–O) mRNA levels of *AstA* (K), *AstC* (L), *Tk* (M), *NPF* (N), and *Dh31*(O) in the gut were measured in *elav-GAL4>UAS-tra^F^*flies and respective genetic controls (*elav-GAL4>+* and *+>UAS-tra^F^*) in females and males. Neuronal Tra expression did not alter *AstA* (female: *p*^GAL4^>0.9999 and *p*^UAS^=0.0534; male: *p*^GAL4^=0.5496 and *p*^UAS^=0.1858), *AstC* (female: *p*^GAL4^=0.5948 and *p*^UAS^=0.0878; male: *p*^GAL4^=0.1745 and *p*^UAS^=0.1745), *Tk* (female: *p*^GAL4^>0.9999 and *p*^UAS^=0.0269; male: *p*^GAL4^>0.9999 and *p*^UAS^=0.2110), *NPF* (female: *p*^GAL4^>0.9999 and *p*^UAS^=0.1158; male: *p*^GAL4^>0.9999 and *p*^UAS^=0.6652), or *Dh31* (female: *p*^GAL4^=0.5442 and *p*^UAS^=0.8086; male: *p*^GAL4^=0.1650 and *p*^UAS^=0.5258) levels in either sex. Sex:genotype interaction: *AstA* (*p*=0.6125), *AstC* (*p*=0.4992), *Tk* (*p*=0.5212), *NPF* (*p*=0.6546), *Dh31* (*p*=0.9566). Data were analyzed by two-way ANOVA with Bonferroni or Tukey’s HSD post-hoc tests as appropriate (aligned rank transform applied for non-parametric data in L and O); *n*=6 biological replicates. (P–T) mRNA levels of *AstA* (P), *AstC* (Q), *Tk* (R), *NPF* (S), and *Dh31* (T) in the head were measured in *elav-GAL4>UAS-tra^F^*flies and respective genetic controls (*elav-GAL4>+* and *+>UAS-tra^F^*) in females and males. Neuronal Tra expression did not alter *AstA* (female: *p*^GAL4^=0.9843 and *p*^UAS^=0.7086; male: *p*^GAL4^=0.0628 and *p*^UAS^=0.9936), *AstC* (female: *p*^GAL4^=0.1253 and *p*^UAS^=0.8540; male: *p*^GAL4^=0.9086 and *p*^UAS^=0.0188), *Tk* (female: *p*^GAL4^=0.6051 and *p*^UAS^=0.9999; male: *p*^GAL4^=0.3600 and *p*^UAS^=0.2760), or *Dh31* (female: *p*^GAL4^>0.9999 and *p*^UAS^>0.9999; male: *p*^GAL4^=0.2918 and *p*^UAS^=0.5990) levels in either sex. *NPF* was reduced in females compared with both controls (*p*^GAL4^=0.0347 and *p*^UAS^=0.0273) but was unchanged in males (*p*^GAL4^=0.0656 and *p*^UAS^=0.6253). Sex:genotype interaction: *AstA* (*p*=0.0171), *AstC* (*p*=0.0198), *Tk* (*p*=0.2324), *NPF* (*p*=0.6872), *Dh31* (*p*=0.4360). Data were analyzed by two-way ANOVA with Tukey’s HSD or Bonferroni post-hoc tests as appropriate (aligned rank transform applied for non-parametric data in P, Q, R, S); *n*=5–6 biological replicates. All data plotted as mean ± SEM. ns indicates not significant with *p*>0.05; * *p*<0.05, ** *p*<0.01, *** *p*<0.001, **** *p*<0.0001. See also Figure S2.

### 2.3. Gut-derived Allatostatin C promotes female fat storage

Given that EE cell-derived AstC, NPF, and Tk regulate fat storage and phenotypes associated with fat storage (e.g., starvation resistance) in single- and mixed-sex animal groups [22,29,44,72], we wanted to assess whether these hormones contribute to the sex difference in fat storage. We used RNAi to knock-down levels of *AstC, Tk* and *NPF* with GAL4 drivers targeting these specific EE populations (*AstC-GAL4, Tk-GAL4,* and *NPF-GAL4*, respectively). Importantly, GAL4 activity for each driver line was restricted to the gut using *R57C10-GAL80*, a validated approach to target only the gut cells that produce these hormones [22,44,45]. For all fat storage data, cell type-specific RNAi was considered to have a significant effect on fat storage only if the experimental genotype (e.g., *tissue-GAL4>UAS-RNAi*) significantly differed from both parental strains (e.g., *tissue-GAL4>+* and *+>UAS-RNAi*) with the same direction of effect. We found gut-specific knock-down of *AstC* (genotype *AstC-GAL4>UAS-AstC-RNAi, R57C10-GAL80*) caused a significant reduction in body fat in females with no effect in males (Figure 3A). Gut-specific knock-down of *Tk* (genotype *Tk-GAL4>UAS-Tk-RNAi, R57C10-GAL80*) similarly showed a trend toward a female-specific decrease in fat storage (*p*^GAL4^=0.1109; *p*^UAS^=0.0118), with no significant effect on male body fat (Figure 3B).

**Figure 3.**
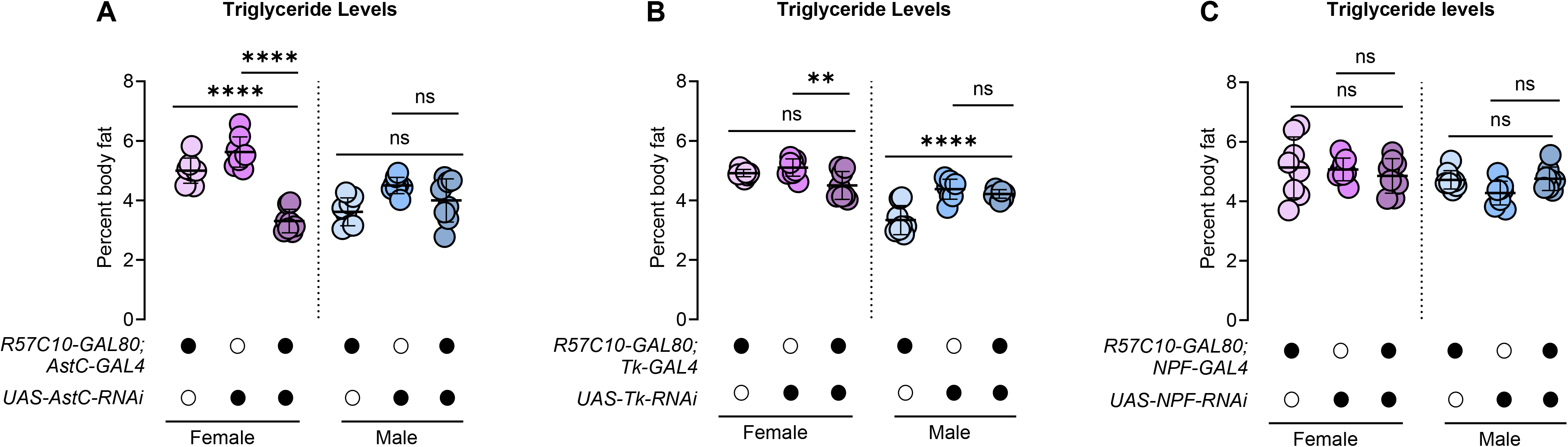
Gut-derived Allatostatin C promotes female fat storage. For all data, cell type-specific RNAi was considered to have a significant effect only if the experimental genotype (e.g., *tissue-GAL4>UAS-RNAi*) significantly differed from both parental strains (e.g., *tissue-GAL4>+* and *+>UAS-RNAi*) with the same direction of effect. (A) Whole-body triglyceride levels were significantly lower in *AstC-GAL4>UAS-AstC-RNAi*,*R57C10-GAL80* females compared with *AstC-GAL4>+,R57C10-GAL80 and +>UAS-AstC-RNAi* control females (*p*^GAL4^<0.0001 and *p*^UAS^<0.0001), an effect that was not observed in males (*p*^GAL4^=0.4527 and *p*^UAS^=0.1056) (sex:genotype interaction *p*<0.0001). Two-way ANOVA followed by Bonferroni post-hoc test; *n*=8 biological replicates. (B) Whole-body triglyceride levels were not significantly different in *Tk-GAL4>UAS-Tk-RNAi*,*R57C10-GAL80* females and males compared with controls (female: *p*^GAL4^=0.1109 and *p*^UAS^=0.0118; male: *p*^GAL4^<0.0001 and *p*^UAS^=0.5704) (sex:genotype interaction *p*<0.0001) though we note a trend toward lower body fat in females. Two-way ANOVA followed by Bonferroni post-hoc test; *n*=8 biological replicates. (C) Whole-body triglyceride levels were not significantly different in *NPF-GAL4>UAS-NPF-RNAi,R57C10-GAL80* females and males compared with controls (female: *p*^GAL4^>0.9999 and *p*^UAS^>0.9999; male: *p*^GAL4^>0.9999 and *p*^UAS^=0.4134) (sex:genotype interaction *p*=0.2890). Two-way ANOVA followed by Bonferroni post-hoc test; *n*=8 biological replicates. All data plotted as mean ± SEM. ns indicates not significant with *p*>0.05; ** *p*<0.01, **** *p*<0.0001.

This suggests a role for gut-derived AstC and a potential role for gut-derived Tk in regulating female body fat, whereas gut-derived AstC or Tk do not play a role in regulating male body fat. In contrast, gut-specific knock-down of *NPF* (*NPF-GAL4>UAS-NPF-RNAi, R57C10-GAL80*) did not significantly alter whole-body fat storage in either males or females (Figure 3C). Together, these data indicate that female-biased expression of AstC in the gut contributes to the regulation of fat storage in females and suggest a possible role for Tk in this process.

### 2.4. Allatostatin C receptor and Tachykinin receptor in neurons promote fat storage in females but not males

Neurons and neuropeptide-producing cells are key cell types upon which AstC, Tk, and NPF act to influence physiology [22,29,32,72,80,81]. Based on our findings with EE cell-specific knock-down of AstC and Tk, we therefore predicted that knock-down of *AstC-R2* and *TkR99D* in these cells may reproduce the reduced fat storage we observed in females with knock-down of EE cell-derived AstC, and the trend toward lower body fat with knock-down of EE cell-derived Tk. To test this, we used *elav*-*GAL4* to knock-down *AstC-R2* and *TkR99D* in post-mitotic neurons and neuropeptide-producing cells.

Importantly, we cannot fully rule out effects of Tk and AstC mediated by other receptors as we did not test these additional receptors [82,83]. In females, knock-down of *AstC-R2* in neurons and neuropeptide-producing cells caused a significant decrease in body fat, with no effect in males (Figure 4A). This reproduced the body fat phenotype caused by knock-down of gut AstC. A female-specific reduction in fat storage was also observed with knock-down of *TkR99D* in neurons and neuropeptide-producing cells (Figure 4B). In line with the lack of body fat effect due to knock-down of gut-derived NPF, we saw no significant change in fat storage in either males or females with knock-down of *NPFR* in neurons and neuropeptide-producing cells (Figure 4C). Together, these data suggest that AstC promotes whole-body fat storage in females via effects on neurons and neuropeptide-producing cells. While EE cell-derived Tk may play a similar role, more work will be needed to test this in future studies.

**Figure 4.**
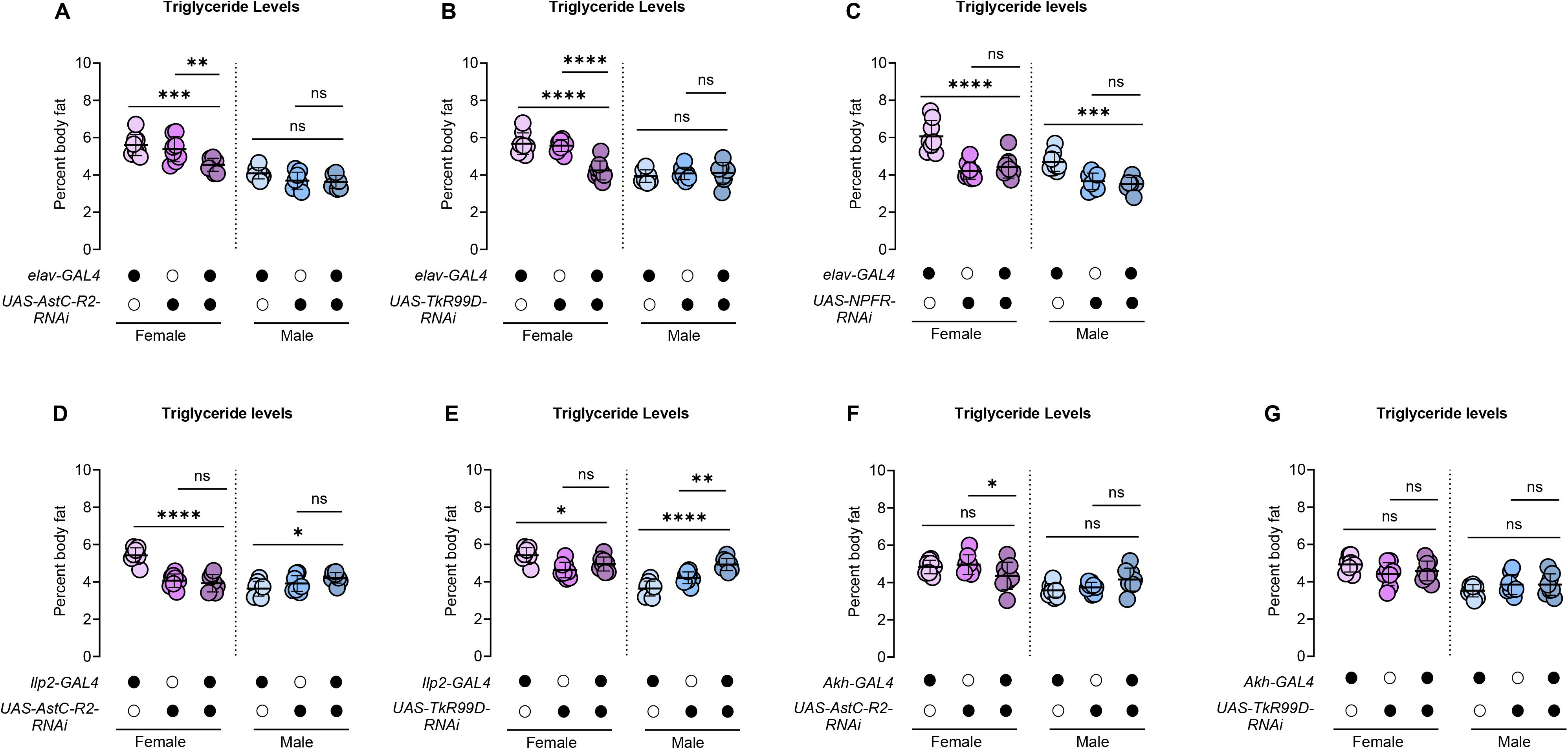
Allatostatin C receptor and Tachykinin receptor in neurons promote fat storage in females but not males. For all data, cell type-specific RNAi was considered to have a significant effect only if the experimental genotype (e.g., *tissue-GAL4>UAS-RNAi*) significantly differed from both parental strains (e.g., *tissue-GAL4>+* and *+>UAS-RNAi*) with the same direction of effect. (A–C) Whole-body triglyceride levels were measured in *elav-GAL4>UAS-RNAi* flies targeting *AstC-R2* (A), *TkR99D* (B), and *NPFR* (C), along with respective genetic controls (*elav-GAL4>+* and *+>UAS-RNAi*) in females and males. Knock-down of *AstC-R2* (female: *p*^GAL4^=0.0001 and *p*^UAS^=0.0013; male: *p*^GAL4^=0.0564 and *p*^UAS^>0.9999; sex:genotype interaction *p*=0.0631) and *TkR99D* (female: *p*^GAL4^<0.0001 and *p*^UAS^<0.0001; male: *p*^GAL4^>0.9999 and *p*^UAS^>0.9999; sex:genotype interaction *p*<0.0001) significantly reduced triglyceride levels in females compared with no significant differences in males. Knock-down of *NPFR* did not change triglyceride levels in either females or males compared with controls (female: *p*^GAL4^<0.0001 and *p*^UAS^>0.9999; male: *p*^GAL4^<0.0001 and *p*^UAS^>0.9999; sex:genotype interaction *p*=0.2470). Data were analyzed by two-way ANOVA followed by Bonferroni post-hoc tests; *n*=8 biological replicates. (D) Whole-body triglyceride levels were not significantly different in *dilp2-GAL4>UAS-AstC-R2-RNAi* females and males compared with controls (female: *p*^GAL4^<0.0001 and *p*^UAS^>0.9999; male: *p*^GAL4^=0.0156 and *p*^UAS^=0.3419; sex:genotype interaction *p*<0.0001). Two-way ANOVA followed by Bonferroni post-hoc test; *n*=8 biological replicates. (E) Whole-body triglyceride levels were not significantly different in *dilp2-GAL4>UAS-TkR99D-RNAi* females compared with controls (*p*^GAL4^=0.0321 and *p*^UAS^=0.0724). Whole-body triglyceride levels were significantly higher in *dilp2-GAL4>UAS-TkR99D-RNAi* males compared with *dilp2-GAL4>+* and *+>UAS-TkR99D-RNAi* control males (*p*^GAL4^<0.0001 and *p*^UAS^=0.0003; sex:genotype interaction *p*<0.0001). Two-way ANOVA followed by Bonferroni post-hoc test; *n*=8 biological replicates. (F) Whole-body triglyceride levels were not significantly different in *Akh-GAL4>UAS-AstC-R2-RNAi* females and males compared with controls (female: *p*^GAL4^=0.3817 and *p*^UAS^=0.0181; male: *p*^GAL4^=0.1229 and *p*^UAS^>0.9999; sex:genotype interaction *p*=0.0241). Two-way ANOVA followed by Bonferroni post-hoc test; *n*=8 biological replicates. (G) Whole-body triglyceride levels were not significantly different in *Akh-GAL4>UAS-TkR99D-RNAi* females and males compared with controls (female: *p*^GAL4^=0.4601 and *p*^UAS^>0.9999; male: *p*^GAL4^>0.9999 and *p*^UAS^=0.8744; sex:genotype interaction *p*=0.0595). Two-way ANOVA followed by Bonferroni post-hoc test; *n*=8 biological replicates. All data plotted as mean ± SEM. ns indicates not significant with *p*>0.05; * *p*<0.05, ** *p*<0.01, *** *p*<0.001, **** *p*<0.0001.

To narrow down the neurons and neuropeptide-producing cells in which *AstC-R2* and *TkR99D* act to mediate their effects on female fat storage, we used cell-type-specific GAL4 drivers to overexpress RNAi transgenes directed at these genes. Given that these gut-derived peptides have been shown to influence metabolic homeostasis and feeding via effects on the insulin-producing cells (IPCs) and the Akh-producing cells (APCs) [19,22,68], we first knocked down *AstC-R2* and *TkR99D* in these cells. We used *dilp2-GAL4* to drive expression of *UAS-Astc-R2-RNAi* and *UAS-TkR99D-RNAi* in the IPCs, and *Akh-GAL4* to drive expression of these transgenes in the APCs. Knock-down of *AstC-R2* in the IPC had no significant effect on fat storage in either males or females compared with sex-matched controls (Figure 4D). IPC-specific knock-down of *TkR99D*, on the other hand, caused a significant increase in whole-body fat storage in males (Figure 4E) with no change in females. In the APC, knock-down of neither receptor altered fat storage in males or females (Figure 4F, 4G). These findings are interesting for several reasons. For example, in males, knock-down of EE cell-derived *Tk* and knock-down of *TkR99D* across neurons had no effect on fat storage, in contrast to the greater fat storage observed with IPC-specific *TkR99D* knock-down. This suggests that Tk derived from outside of the gut, and likely in the head, regulates fat storage via effects on *TkR99D* in the IPC. Further studies will be needed to understand why IPC but not pan-neuronal knock-down of TkR99D causes an effect on male body fat. Possible explanations include greater knock-down in the IPC using *Dilp2-GAL4,* or that Tk mediates opposing effects on body fat via effects on *TkR99D* in multiple neuron groups.

## 3. DISCUSSION

EE cell-derived hormones regulate body fat in single- and mixed-sex animal groups; however, it has been unclear whether the regulation and function of these peptides differ between the sexes. The goal of our study was to perform a detailed comparison of EE cell-derived hormones between the sexes, and to test if these hormones contribute to the sex difference in fat storage. Our assessment revealed profound female-biased expression of EE cell-expressed hormones within the gut. For one hormone, AstC, this differential expression was physiologically significant, as we showed that EE cell-derived AstC promote female fat storage. Interestingly, these effects were not mediated by the IPC or APC, cells that we have previously shown contribute to the sex difference in fat storage. Taken together, our data provide additional insight into the highly complex mechanism(s) by which unmated female flies achieve higher fat storage than male flies (Figure 5).

**Figure 5.**
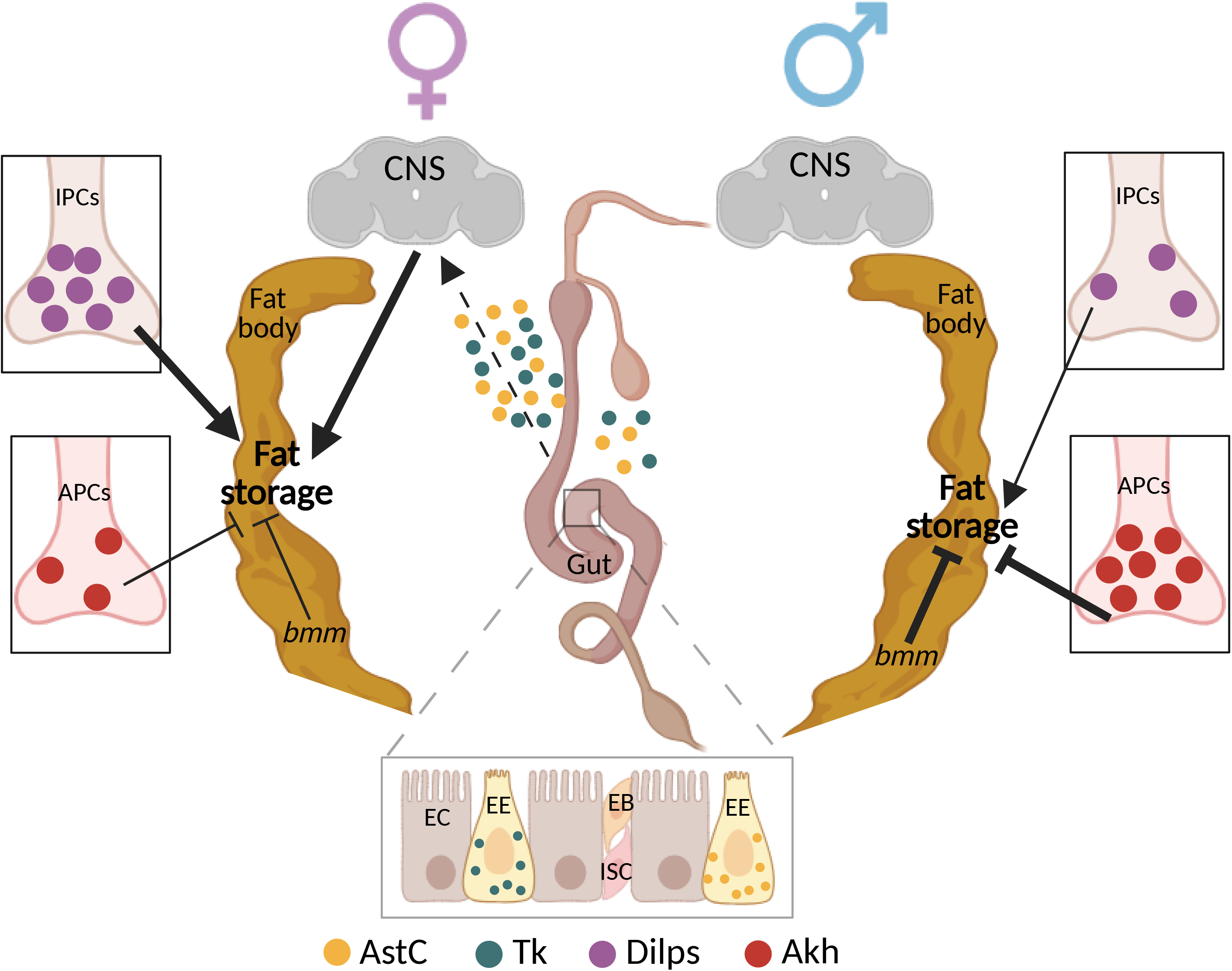
Model of sex differences in fat storage. Schematic representation of complex mechanisms that promote higher fat storage in *Drosophila* females. Profound female-biased EE cell-derived Tk and AstC promote female fat storage, an effect mediated by neurons but independent of the IPC or APC. EE cell-derived factors therefore act alongside a greater insulin/insulin-like growth factor signaling pathway activity, lower Adipokinetic hormone (Akh) signaling, and lower expression of triglyceride lipase *brummer* (*bmm*) to specify higher adiposity in female flies. In males, we detected no contribution of gut-derived factors to fat storage, which we previously showed is kept at a lower level than in females by higher *bmm* expression and Akh signaling. Thus, multiple complex mechanism specify higher fat storage in unmated female flies compared with males. IPCs: insulin-producing cells; APCs: adipokinetic hormone-producing cells; CNS: central nervous system; *bmm*: *brummer*; EC: enterocyte; EB: enteroblast; EE: enteroendocrine; ISC: intestinal stem cells; AstC: Allatostatin C; Tk: Tachykinin; Dilps: *Drosophila* insulin-like peptides; Akh: Adipokinetic hormones. Created in BioRender. Rideout, E. (https://BioRender.com/9z1gko5) is licensed under CC BY 4.0.

While it was not the main goal of our study, our survey of EE cell-expressed hormones in *Drosophila* revealed that the sex bias in expression was not uniform across tissues. In the gut, mRNA levels of *AstC*, *Tk*, and *NPF* were higher in females than in males. In the brain, mRNA levels of these hormones showed a significant male bias, in line with data from previous reports on Tk [57] and NPF [64]. While it remains unclear whether the tissue-specific sex bias in expression is physiologically significant, peptides derived from the gut and the brain have been shown to mediate distinct effects on physiology and/or behavior. For example, EE cell-derived AstC regulates energy homeostasis and food-seeking behaviors in adult females [22], whereas neuron-derived AstC is involved in regulating locomotion in adult males [60] and the circadian regulation of oogenesis in adult females [70]. Neuron-derived Tk similarly regulates locomotion [65,69], food consumption [68], Dilp secretion [19], and aggression [57], whereas gut-derived Tk regulates intestinal lipogenesis [72] and stem cell divisions in the midgut [84–86]. Supporting a potential sex-specific role for peptides derived from different anatomical sites in regulating physiology, gut-derived AstC stimulates fat breakdown during starvation through the Akh pathway in mated females with no effect in males [22]. Future studies are therefore needed to determine whether there are sex differences in whether the effects of EE cell-derived hormones are primarily mediated by local or systemic mechanisms.

Another important task for future studies will be to elucidate how sex differences in neuropeptide expression are established. The first step in understanding these mechanisms will be to determine which factors specify the sex bias in neuropeptide mRNA levels. Because our data shows that sex determination gene *tra* does not regulate the sex bias in neuropeptide expression in either the brain or the gut, the role of other factors that influence sexual identity and sexual differentiation must be assessed. One strong candidate is the steroid hormone ecdysone, as virgin females have higher ecdysone titers than males [87–89]. Ecdysone plays a role in regulating sexual differentiation and development [5,90,91], and contributes to male-female differences in multiple aspects of intestinal physiology (e.g., intestinal stem cell proliferation) [46,50] and brain development [92,93]. Another candidate is juvenile hormone, which has been shown to regulate sexual maturation in *Drosophila* and other insects [94–100]. While it remains unclear whether juvenile hormone titers differ between virgin males and females, juvenile hormone regulates many aspects of gut physiology in mated females (e.g., intestinal lipid accumulation, intestinal stem cell proliferation) [54,101] and influences brain development [102]. Other than hormones, it is possible that sex determination gene *Sex-lethal* plays a role in regulating the sex difference in mRNA levels of EE cell-derived hormones, as *tra*-independent effects of *Sex-lethal* have been described in the brain [103]. While sex differences in expression of EE cell-derived hormones does not involve *tra*, and is therefore unlikely to involve known *tra* targets such as *fruitless* [104,105], without further experiments we cannot fully rule out these additional sex determination pathway members.

In parallel to identifying the factor(s) responsible for establishing the sex difference in EE cell-expressed hormones, it will be important to reveal the cellular basis for this differential expression. For example, a sex difference in the number of EE cells and neuropeptide-expressing cells in the brain could explain the differences in expression. Supporting this, gut length, overall brain size, and neuron number have been shown to differ between males and females [34,48,106–113]. In the gut, the difference in length is at least partially due to a sex difference in the proliferation of intestinal stem cells [46,48,49], which undergo asymmetric divisions and subsequent differentiation to generate all gut cell types including EE cells [34,42,114–118].

In the brain, males and females differ in the number of neurons found within many identified clusters [106,107,109–113,119], including the cells that produce NPF [64] and Tk [57]. Differences in neuron number have been primarily attributed to sex-specific programmed cell death [112,119–122]; however, sex differences in neuroblast cell death and/or proliferation may also play a role [111,123–125].

Beyond the effects of cell number, sex differences in EE cell-derived hormone mRNA levels may also be due to differential gene and/or protein expression of these factors, which have been reported for other peptide hormones [7,18,64]. Because sex differences in the activity of peptide hormone-producing cells have also been previously described [7], it is clear that a detailed examination of sex differences in EE cells, and more generally in neuropeptide-producing cells, is needed to gain a comprehensive picture of how these cells differ between males and females. Benefits of such a detailed study include gaining insight into potential mechanisms underlying sex differences in other aspects of physiology and behavior. For example, EE cells regulate ISC homeostasis [84,126], and EE cell-derived hormones act locally and systemically to regulate appetite, food ingestion, food digestion, gut motility, and immune responses [35,81,127,128]. Importantly, male-female differences in many of these phenotypes have been reported [52,53,129–132].

Overall, our findings identify EE cell-derived hormones AstC as an important factor that promotes higher fat storage in *Drosophila* adult virgin females. This builds on a recent paper identifying a key role for IIS in promoting higher levels of fat storage in unmated females but not males [18], advancing knowledge of the factors that establish an optimal level of stored fat in each sex.

## 4. MATERIALS AND METHODS

### 4.1. Fly strains

The following fly strains from the Bloomington *Drosophila* Stock Center were used: *w^1118^* (RRID:BDSC_3605), *voila-GAL4* (RRID:BDSC_80572), *dilp2-GAL4* (IPCs) (RRID:BDSC_37516), *UAS-tra^F^* (RRID:BDSC_4590), *UAS-NPF-RNAi* (RRID:BDSC_27237), *UAS-TkR99D-RNAi* (RRID:BDSC_27513), *UAS-AstC-RNAi* (RRID:BDSC_25868), *UAS-NPFR-RNAi* (RRID:BDSC_25939), *UAS-AstC-R2-RNAi* (RRID:BDSC_36888), *UAS-Tk-RNAi* (RRID:BDSC_25800), *elav-GAL4* (RRID:BDSC_458). We obtained *R57C10-GAL80; NPF-GAL4, R57C10-GAL80; Tk-GAL4 and R57C10-GAL80; AstC-GAL4* as kind gifts from Dr. Kim Rewitz at the University of Copenhagen, and *Akh-GAL4* was a kind gift from Dr. Mike Gordon at The University of British Columbia. We acknowledge FlyBase as an essential resource providing genetic, genomic, and functional data and tools that supported this study [133].

### 4.2. Fly husbandry

Fly media was prepared with the following ingredients: 20.5 g/L sucrose, 70.9 g/L D-glucose, 48.5 g/L cornmeal, 45.3 g/L yeast, 4.55 g/L agar, 0.5 g/L CaCl_2_•2H_2_O, 0.5 g/L MgSO_4_•7H_2_O, 11.77 mL/L acid mix (propionic acid/phosphoric acid). For all experiments, we allowed female flies to lay eggs on grape juice agar plates for 12 hr. At 24 hr after egg laying, 50 larvae were picked into vials containing 10 mL of food and reared at 22°C. Males and females were distinguished by the presence of sex combs in the late pupal period and placed into single-sex vials to eclose. After eclosion, adult flies were maintained at a density of twenty flies per vial in single-sex groups. Unless otherwise stated, all experiments used 5- to 7-day-old unmated flies. We used unmated flies to identify genetic factors that regulate the sex difference in body fat; mated females were not used to avoid mating-induced changes in physiology mediated by additional factors (*e.g*., Sex-peptide) and behavioral changes due to altered food preferences.

### 4.3. Adult weight

Groups of 10 flies were placed in pre-weighed 1.5 ml microcentrifuge tubes (Diamed Lab Supplies, DIATEC610-2550) and weighed on an analytical balance (Mettler-Toledo, ME104).

### 4.4. Whole-body triglyceride measurements

Triglyceride is the main form of stored fat in the body, with very little in the circulation [134]. We therefore refer to whole-body triglyceride as ‘fat storage’ or ‘body fat’. One biological replicate consisted of five flies. Flies were collected in a 1.5 ml tube and homogenized in 350 μl of 0.1% Tween (Amresco, 0777-1L) in 1X phosphate-buffered saline (PBS; Sigma-Aldrich, P5493) using 50 μl of glass beads (Sigma-Aldrich, Z250473) that were agitated at 8 m/s for 5 s (OMNI International BeadRuptor 24).

Triglyceride concentration was measured using the Stanbio Triglyceride Liquid Reagent (FT7610, BD386a/d, BD386b) according to the manufacturer’s instructions and as described previously [13,18] with minor modifications. Briefly, 10 μl of either homogenate or triglyceride standard (FT7610) was added to 190 μl of activated triglyceride reagent (Enzymatic Triglyceride Reagent, BD386a/d; Triglyceride Activator, Cat. No. BD386b) in a 96-well plate. After a 15 min incubation at room temperature, the absorbance was read at 540 nm (Thermo Scientific – Multiskan FC Microplate Photometer).

### 4.5. RNA extraction, cDNA synthesis, and Quantitative real-time PCR (qPCR)

One biological replicate consisted of 3-5 adult fly guts, or 10 adult fly heads, or 5 whole-body adult flies. Samples were homogenized in 500 μl Trizol (Thermo Fisher Scientific; 15596018). Chloroform was added to Trizol to separate the mixture into aqueous and organic phases, and isopropanol was added to the aqueous phase (in a fresh tube) to precipitate the RNA. RNA was resuspended in either 20-25 μl (for guts and heads) or 200 μl (for whole-body) of molecular biology grade water (Corning, 46-000-CV). RNA was stored at -80°C until use. Each experiment contained 5-10 biological replicates per sex and per genotype; each experiment was repeated twice.

For genomic DNA elimination and cDNA synthesis, an equal amount of RNA per reaction was DNase-treated and reverse transcribed according to manufacturer’s instructions using the QuantiTect Reverse Transcription Kit (Qiagen, 205314). Relative mRNA transcript levels were quantified using qPCR as described previously [6]. Data were normalized to the average fold change of *Actin5C* and β*-tubulin*. For a full primer list, refer to Table 1.

### 4.6. Statistical analysis

Statistical analyses and data presentation were completed using GraphPad Prism 9 (GraphPad Software, San Diego, CA, USA). All data were tested for normality using the Shapiro-Wilk test. Normally-distributed data were subjected to parametric tests as appropriate, including Student’s *t*-test and two-way ANOVA followed by Bonferroni post-hoc test. For non-normally distributed data, we used the Mann-Whitney test. For two-way ANOVA involving data that do not satisfy the normality assumption, aligned rank transformation was first applied using the art() function from the ARTool R package [135]. Then, ANOVA was performed on the transformed data with the base R anova() function. Finally, the art.con() function from the ARTool package was used to extract the main as well as the interaction effects. Default parameters were used in each step of the analysis. For all statistical analyses, differences were considered significant if *p*<0.05.

## Supporting information

Table 1-primer list

Source Data 1. Excel file containing raw data with calculations

Source Data 2. Excel file conatining statistics for all data

## CRediT authorship contribution statement

**Puja Biswas:** Conceptualization, Data curation, Formal analysis, Investigation, Methodology, Visualization, Writing – original draft and editing.

**Elizabeth J. Rideout:** Conceptualization, Funding acquisition, Methodology, Resources, Supervision, Writing – original draft, Writing – review and editing.

## Acknowledgments

The authors thank FlyBase, which is supported by a grant from the National Human Genome Research Institute at the U.S. National Institutes of Health (U41 HG000739) and by the British Medical Research Council (MR/N030117/1). Stocks obtained from the Bloomington *Drosophila* Stock Center (NIHP40OD018537) were used in this study. The authors thank the TRiP at Harvard Medical School (NIH/NIGMS R01-GM084947) for providing transgenic RNAi fly stocks and/or plasmid vectors used in this study. The funders had no role in study design, data collection and analysis, decision to publish, or preparation of the manuscript. We thank members of the Rideout lab for valuable feedback. We acknowledge that our research takes place on the traditional, ancestral, and unceded territory of the Musqueam people; a privilege for which we are grateful.

## Figure supplements

Figure 1-figure supplement 1

Figure 2-figure supplement 2

**Table**

Table 1-primer list

## Data availability

Source Data 1. Excel file containing raw data with calculations

Source Data 2. Excel file containing statistics for all data

