## Supplementary figures and images for "Sex-biased expression of enteroendocrine cell-derived hormones contributes to higher fat storage in *Drosophila* females"

### Figure 1-figure supplement 1

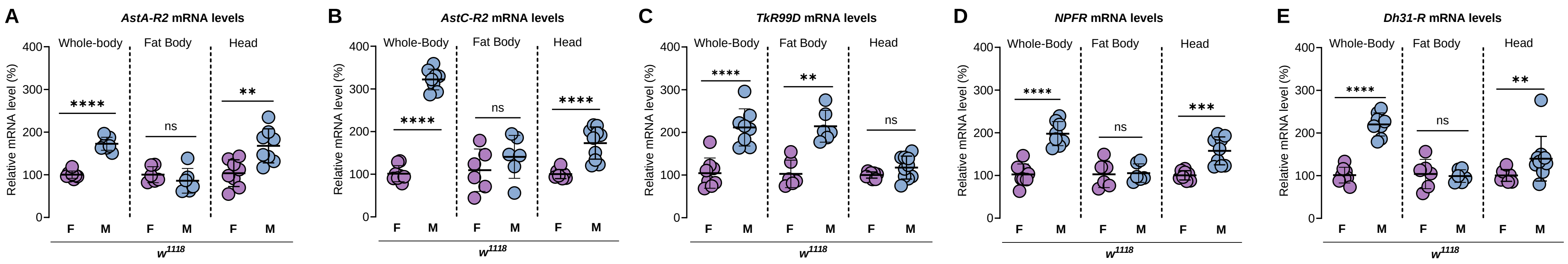

### Figure 2-figure supplement 2

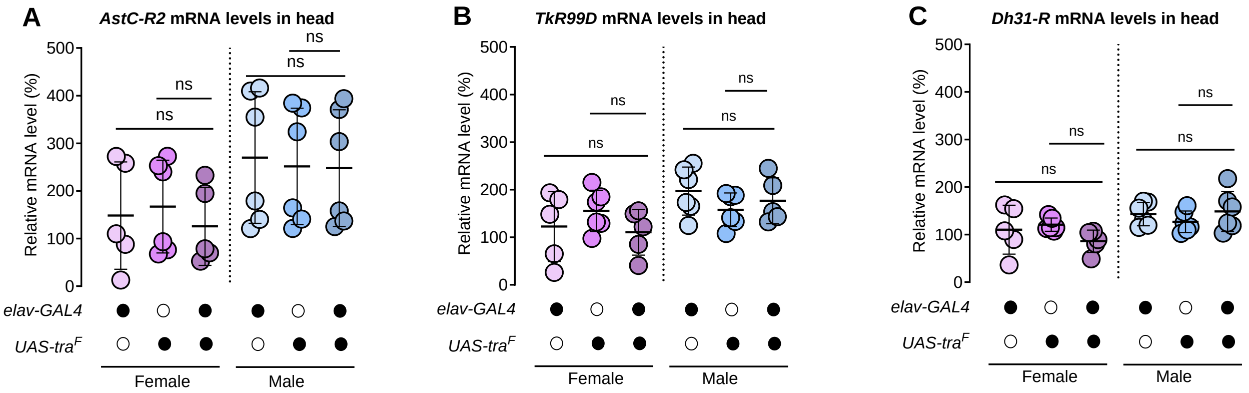
