## Supplementary material for "Sex-biased expression of enteroendocrine cell-derived hormones contributes to higher fat storage in *Drosophila* females": Table 1-primer list

List of primers

| Gene name | Forward (5ʹ-3ʹ) | Reverse (5ʹ-3ʹ) |
| --- | --- | --- |
| Beta tubulin | ATCATCACACACGGACAGGA | GAGCTGGATGATGGGGAGTA |
| Actin5C | TTGTCTGGGCAAGAGGATCAG | ACCACTCGCACTTGCACTTTC |
| AstA | TTTAGTCCGCGGAACCTCTG | GCTGCTGCTACTGAGCGAAT |
| AstC | TACGGCCTACTCCTCACCC | GCTGGCATATCGTAGCCACC |
| Tk | TGGCAAGAAGAGCGATCTGG | CCTACTCGAAAAGTGCTGGC |
| NPF | GGCTGATGCCTACAAGTTCCT | CTCATTAAAACCGCGAGCAAATTC |
| Dh31 | TCTCAAAGCGGTGCAGTCAG | TGCGGCTGTCTCCCTTTTTC |
| AstA-R2 | CGAACACCCTCACCAAGCTA | GGAGCAGTTAACGGCCTTGT |
| AstC-R2 | ACTGAATCTGGCTATCGCGG | TGCTCACCATGTAGGCCTTG |
| TkR99D | GATGAATTCGCGCTTTCGCT | ATTCGATGCGACTTGGGTGA |
| NPFR | TTTGCATGCTCCAAACGTCG | CAGCCAGAGTGTTTCCCGAT |
| Dh31-R | CCACTCAGGTCTCGTTCTTTTG | CTGCTGATCCGTGGACAACT |
