## Supplementary material for "Sex-biased expression of enteroendocrine cell-derived hormones contributes to higher fat storage in *Drosophila* females": figure supplement 1 legends

**Figure 1-figure supplement 1 – Sex differences in mRNA levels of receptors for EE cell-derived hormones.**

(A) mRNA levels of *AstA-R2* in whole-body (*p*<0.0001; Student’s *t*-test) and in head (*p*=0.0031; Student’s *t*-test) were significantly higher in 5-day-old *w^1118^* males compared to females; no sex difference was observed in fat body (*p*=0.3329; Student’s *t*-test). n=6-8 biological replicates.

(B) mRNA levels of *AstC-R2* in whole-body (*p*<0.0001; Student’s *t*-test) and in head (*p*<0.0001; Student’s *t*-test) were significantly higher in 5-day-old *w^1118^* males compared to females, no sex difference was observed in fat body (*p*=0.3036; Student’s *t*-test). n=6-10 biological replicates.

(C) mRNA levels of *TkR99D* in whole-body (*p*<0.0001; Student’s *t*-test) and in fat body (*p*=0.0022; Mann-Whitney test) were significantly higher in 5-day-old *w^1118^* males compared to females, no sex difference was observed in heads (*p*=0.0946; Student’s *t*-test). n=6-10 biological replicates.

(D) mRNA levels of *NPFR* in whole-body (*p*<0.0001; Student’s *t*-test) and in head (*p*=0.0003; Student’s *t*-test) were significantly higher in 5-day-old *w^1118^* males compared to females, no sex difference was observed in fat body (*p*=0.8669; Student’s *t*-test). n=6-9 biological replicates.

(E) mRNA levels of *Dh31-R* in whole-body (*p*<0.0001; Student’s *t*-test) and in head (*p*=0.0085; Mann-Whitney test) were significantly higher in 5-day-old *w^1118^* males compared to females, no sex difference was observed in fat body (*p*=0.7585; Student’s *t*-test). n=6-10 biological replicates.

All data plotted as mean ± SEM. ns indicates not significant with *p*>0.05; ** *p*<0.01, *** *p*<0.001, **** *p*<0.0001.
