## Supplementary material for "Sex-biased expression of enteroendocrine cell-derived hormones contributes to higher fat storage in *Drosophila* females": figure supplement 2 legends

**Figure 2-figure supplement 2 – Sex determination gene *transformer* does not regulate mRNA levels of receptors for EE cell-derived hormones.**

(A) mRNA levels of *AstC-R2* in head were not significantly different in *elav-GAL4>UAS-tra^F^* females and males compared with controls (female: *p*^GAL4^=0.9977 and *p*^UAS^=0.9838; male: *p*^GAL4^=0.9998 and *p*^UAS^=1.0) (sex:genotype interaction *p*=0.8735). Two-way ANOVA followed by Tukey HSD on data processed using the aligned rank transform for non-parametric data; n=5-6 biological replicates.

(B) mRNA levels of *TkR99D* in head were not significantly different in *elav-GAL4>UAS-tra^F^* females and males compared with controls (female: *p*^GAL4^>0.9999 and *p*^UAS^=0.4552; male: *p*^GAL4^>0.9999 and *p*^UAS^>0.9999) (sex:genotype interaction *p*=0.1903). Two-way ANOVA followed by Bonferroni post-hoc test; n=5-6 biological replicates.

(C) mRNA levels of *Dh31-R* in head were not significantly different in *elav-GAL4>UAS-tra^F^* females and males compared with controls (female: *p*^GAL4^=0.7154 and *p*^UAS^=0.2409; male: *p*^GAL4^>0.9999 and *p*^UAS^=0.7312) (sex:genotype interaction *p*=0.1210). Two-way ANOVA followed by Bonferroni post-hoc test; n=5-6 biological replicates.

All data plotted as mean ± SEM. ns indicates not significant with *p*>0.05.
